# Characterisation and structure determination of a llama-derived nanobody targeting the J-base binding protein 1

**DOI:** 10.1101/340778

**Authors:** Bart Van Beusekom, Tatjana Heidebrecht, Athanassios Adamopoulos, Alexander Fish, Els Pardon, Jan Steyaert, Robbie P. Joosten, Anastassis Perrakis

**Affiliations:** Biochemistry, Netherlands Cancer Institute, Plesmanlaan 121, Amsterdam, 1066 CX, The Netherlands; VIB Center for Structural Biology, Vrije Universiteit Brussel, Pleinlaan 2, Brussels, 1050, Belgium; Structural Biology Brussels, Vrije Universiteit Brussel, Pleinlaan 2, Brussel, 1050, Belgium

**Keywords:** nanobody, JBP1, llama, immune system

## Abstract

The J-base Binding Protein 1 (JBP1) contributes to biosynthesis and maintenance of base J (*β*-D-glucosyl-hydroxymethyluracil), a modification of thymidine confined to some protozoa. Camelid (llama) single domain antibody fragments (nanobodies) targeting JBP1 were produced for use as crystallization chaperones. Surface plasmon resonance (SPR) screening identified Nb6 as a strong binder, recognising JBP1 with a 1:1 stoichiometry and high affinity (k_D_=30nM). Crystallisation trials of JBP1 in complex with Nb6, yielded crystals diffracting to 1.47Å resolution.

However, the asymmetric unit dimensions and molecular replacement with a nanobody structure, clearly showed that the crystals of the expected complex with JBP1 were of the nanobody alone. Nb6 crystallizes in spacegroup P31 with two molecules in the asymmetric unit; its crystal structure was refined to a final resolution of 1.64Å. Ensemble refinement suggests that on the ligand-free state one of the complementarity determining regions (CDRs) is flexible while the other two adopt well-defined conformations.

**Synopsis:** A camelid single domain antibody fragment (nanobody) is shown to have high affinity towards its recognition target, the J-base binding protein 1 (JBP1). The serendipitous crystallisation of this nanobody alone, and its crystal structure solution and refinement to 1.64Å resolution are described. Ensemble refinement suggests that on the ligand-free state one of the complementarity determining regions (CDRs) is flexible while the other two adopt well-defined conformations.

## 1. Introduction

In kinetoplastid flagellates such as Trypanosoma, Leishmania and Crithidia, one percent of thymidine is replaced by a modified base called base-J (*β*-D-glucosyl-hydroxymethyluracil) (Gommers-Ampt *et al.*, 1993). In Leishmania, 99 percent of J is located in telomeric repeats (van Leeuwen *et al.*, 1996; van Leeuwen *et al.*, 1997; Genest *et al.*, 2007).

The remaining one percent resides in internal chromosomal positions (iJ). In Trypanosomes this iJ is associated with transcription initiation sites (Cliffe *et al.*, 2010), while in Leishmania iJ is associated with transcription termination (van Luenen *et al.*, 2012).

A two-step mechanism is responsible for the biosynthesis of base-J. In the first step, hydroxymethyluracil (hmU) is formed by hydroxylation of the 5-methyl group of specific thymines in the genome of kinetoplastida. Second, a glucose molecule is transferred to hmU, creating base-J (Bullard *et al.*, 2014; Iyer *et al.*, 2013; Sekar *et al.*, 2014). Two proteins are responsible for the first step, J-DNA binding protein 1 and 2 (JBP1, JBP2) that both contain an N-terminal hydroxylase domain. This hydroxylation reaction, which is dependent on the presence of Fe(II) and 2-oxoglutarate (2OG) (Cliffe *et al.*, 2012), converting T to hMU, provided the inspiration for suggesting and proving the mechanism for the conversion of methylcytosin (mC) to hydroxymethylcytsine (hmC) in humans by the TET proteins (Tahiliani *et al.*, 2009; Ito *et al.*, 2010). While the structure of the hydroxylase domain of TET proteins has been resolved (Hu *et al.*, 2013) the structure of the JBPs hydroxylase domain remains elusive.

JBP1 specifically recognises J-base containing DNA (J-DNA), through a short ~150 residue autonomous folding unit in the middle of JBP1: the J-DNA binding domain (JDBD). Equilibrium kinetics experiments revealed that JDBD has a high affinity for base-J (10nM) and remarkable specificity for J-DNA over normal DNA (~10,000 fold). Pre-steady state kinetics experiments together with small angle X-ray and neutron scattering (SANS and SAXS) experiments, suggest that JBP1 has increased flexibility and undergoes a conformational change upon recognition and binding of J-DNA (Heidebrecht *et al.*, 2011; Heidebrecht *et al.*, 2012). This flexible nature of JBP1 has proven to be the most likely bottleneck that has prevented us to crystallise JBP1 over countless efforts.

To decrease JBP1 flexibility, we decided to use nanobodies as crystallization chaperones. In the early 1990s it was discovered (Hamers-Casterman *et al.*, 1993) that camelids possess a type of immunoglobulin (Ig) with only one heavy chain and and three Ig domains, compared to the classical Ig composed of a light chain with two Ig domains and a heavy chain with four Ig domains. Nanobodies are the recombinant variable domains of camelid heavy-chain antibodies, produced by immunization of llamas, and have been a valuable tool in structural biology. They are often used as crystallization chaperones, which will bind with high affinity to target molecules decreasing their flexibility and providing the necessary crystal contacts (Korotkov *et al.*, 2009; Rasmussen *et al.*, 2007).

We have raised nanobodies that bind specifically to JBP1. Here, we present the characterisation of nanobody-6 (Nb6) which binds JBP1 with high affinity, and its serendipitous and somewhat unfortunate crystallisation in the absence of JBP1. This is therefore also a cautionary tale on nanobody-aided crystallisation: although it rarely happens, excellent nanobodies against their targets can crystallise alone. The structure of Nb6 has been refined, and we present the ensemble refinement which showed that only one of the complementarity determining regions (CDRs) is flexible while the other two adopt well-defined conformations.

## 2. Materials and methods

### 2.1. Macromolecule production

Constructs encoding for nanobodies were transformed in *Escherichia coli* (WK6 strain) (Table 1) and were grown and purified as described (Pardon *et al.*, 2014). In short: cells were grown in teriffic broth (TB) media supplemented with 0.1% glucose, 2 mM MgCl_2_ and 0.1 mg/mL ampicillin at 37°C, until they reach an optical density of ~ 1.2. Addition of 1mM IPTG induced nanobody overexpression over a period of 18 hours at 25 °C. Cells were collected by centrifugation at 4000 g for 15 min. Osmotic shock was performed by adding 15 mL of TES buffer (200 mM Tris/HCl pH 8.0, 500 mM sucrose, 0.5 mM EDTA) to the pellet and incubating at 4 °C with agitation overnight. Next, 30 mL of a 4x diluted TES buffer was added to the culture followed by incubation on ice for 60 min. The culture was centrifuged at 9000 g for 40 min and the supernatant was subjected to immobilized metal affinity chromatography using Ni-chelating sepharose beads. Binding to the beads was performed for 60 min at 4°C followed by two washing steps, using 10 mL Wash Buffer 1 (50 mM Na phosphate pH 7, 1M NaCl) and 30 mL Wash Buffer 2 (50 mM Na phosphate pH 6, 1M NaCl). Elution was performed in 5 mL of Elution-Buffer (50 mM Na phosphate pH 7, 150 mM NaCl, 300 mM Imidazole). Imidazole was removed by dialysis and fractions containing nanobodies, were analyzed using SDS-PAGE.

**Table 1.**
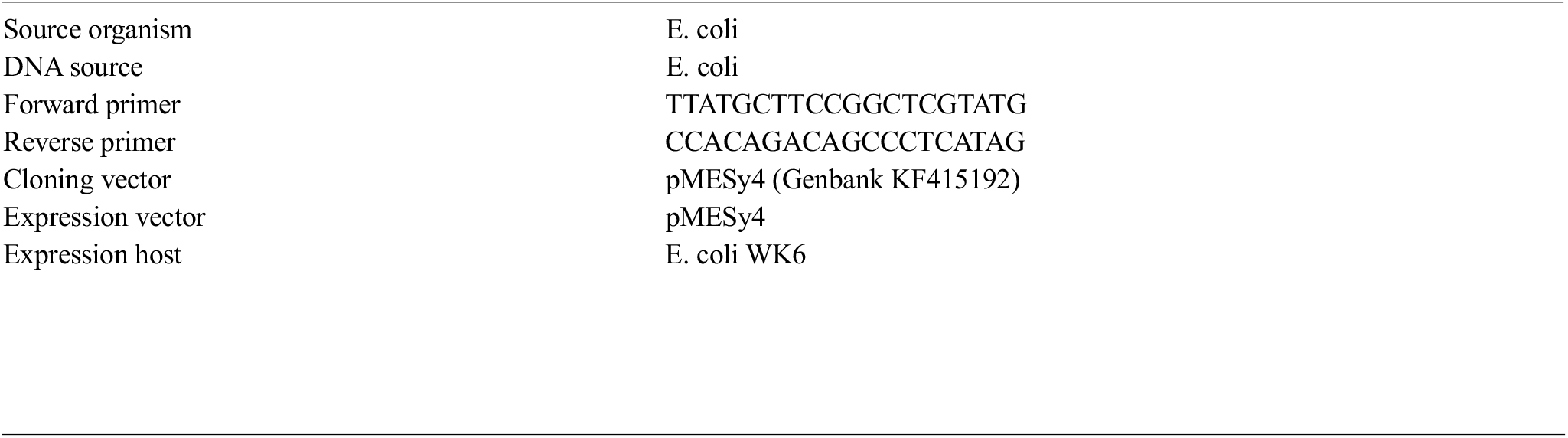
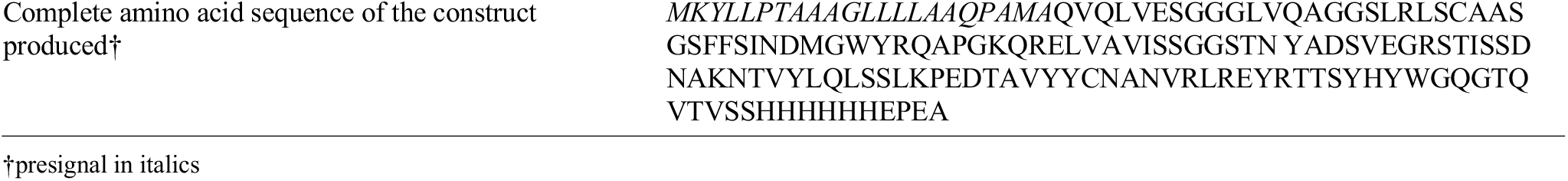

JBP1 with a strep-tag was purified as previously described (van Luenen *et al.*, 2012).

### 2.2. Surface plasmon resonance

The binding affinity towards JBP1 was determined by surface plasmon resonance using a Biacore T200 (Life Sciences General Electric). JBP1 containing a strep tag was immobilized on a Series S Sensor Chip SA (Life Sciences General Electric). The experiment was performed in a buffer consisting of 20 mM HEPES/HCl (pH 7.5), 140 mM NaCl, 0.05% Tween-20, 1mM TCEP, and 1mg/mL BSA. Increasing concentration of Nb6 was injected consecutively over the chip.

The data was analyzed using the Biacore T200 Evaluation software and GraphPad Prism 7 (GraphPad Software Inc.). Equilibrium dissociation constant (KD) was calculated using equation RU=Bmax*Conc/(Conc+Kd).

### 2.3. Crystallization, data collection and processing

The protein-nanobody complex was set up in a 96-well 3-drop Swissci plates sitting-drop format (Table 2). The drops consisted of 100 nL protein/nanobody complex with 100 nL of reservoir. After two days crystals appeared at 277 K in 25% w/v PEG3350 and 0.3 M citric acid (pH 3.5). For vitrifying the crystals for data collection 30% v/v glycerol was added as cryoprotectant. Diffraction data were collected at 100K at the European Synchrotron Radiation Facility on the MASSIF beamline (Bowler *et al.*, 2015; Svensson *et al.*, 2015). Details of data collection and processing are given in Table 3.

**Table 2.**
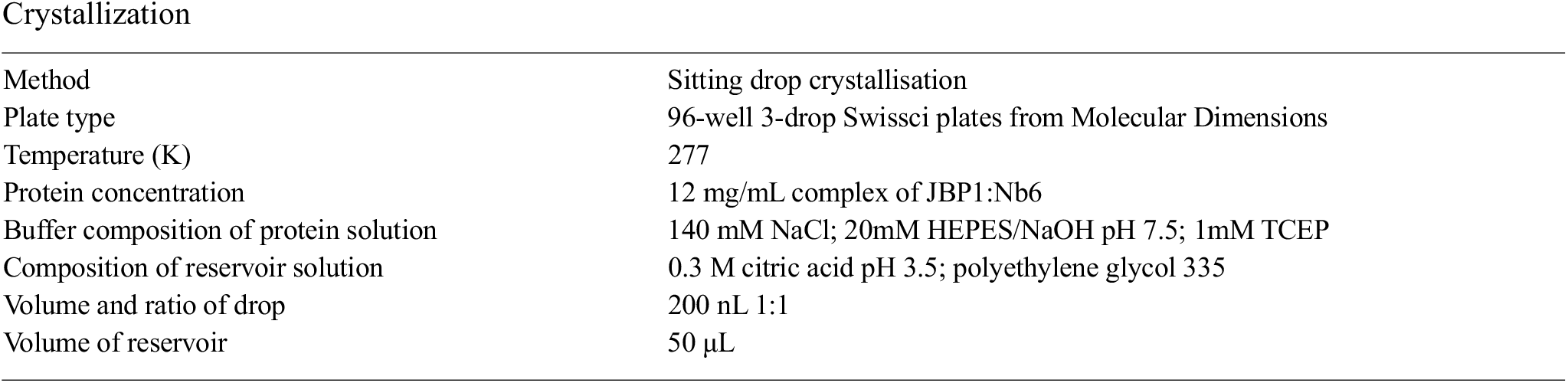
Crystallization

**Table 3.** Data collection and processing

Data collection and processing
|  |  |
| --- | --- |
| 170 Diffraction source | ESRF beamline massif-1 |
| 171 Wavelength (Å) | 0.96600 |
| 172 Temperature (K) | 100 |
| 173 Detector | Dectris PILATUS3 2M pixel |
| 174 Crystal-detector distance (mm) | 171.65 |
| 175 Rotation range per image (°) | 0.20 |
| 176 Total rotation range (°) | 180 |
| 177 Exposure time per image (s) | 0.1 |
| 178 Space group | $P3_1$ |
| 179 $a, b, c$ (Å) | 57.68, 57.68, 64.75 |
| 180 $\alpha, \beta, \gamma$ (°) | 90, 90, 120 |
| 181 Mosaicity (°) | 0.07 |
| 182 Resolution range (Å) | 49.95–1.47 (1.52–1.47) |
| 183 Total No. of reflections | 210220 |
| 184 No. of unique reflections | 40857 |
| 185 Completeness (%) | 99.7 (98.4) |
| 186 Redundancy | 5.100 (5.10) |
| 187 $\langle I/\sigma(I) \rangle$ | 14.3 (1.1) |
| 188 $R_{\text{r.i.m.}}$ | 0.057 (1.541) |
| 189 Overall $B$ factor from Wilson plot (Å<sup>2</sup>) | 23.2 |

**Table 4.**
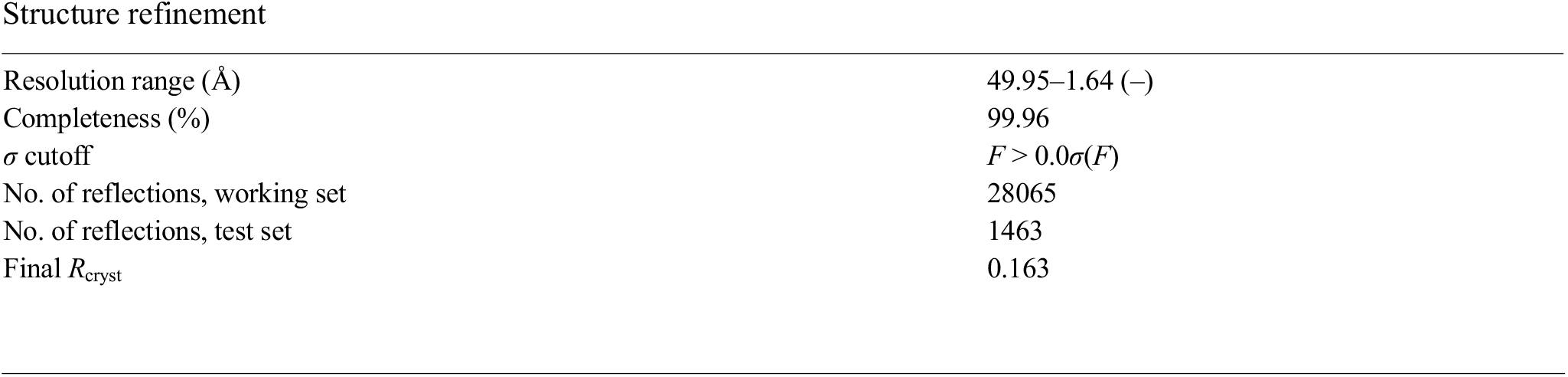
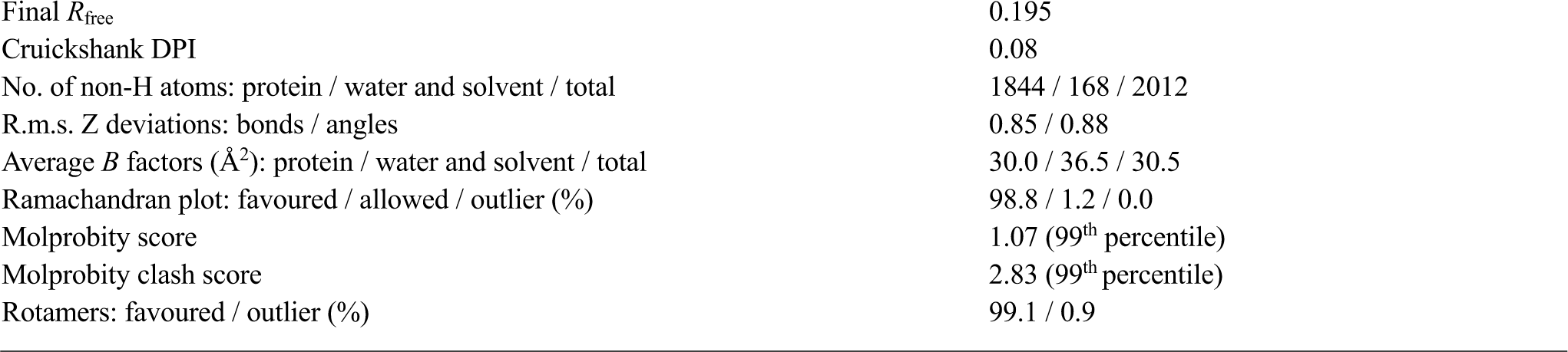
Structure refinement

Structure refinement
|  |  |
| --- | --- |
| 192 Resolution range (Å) | 49.95–1.64 (–) |
| 193 Completeness (%) | 99.96 |
| 194 $\sigma$ cutoff | $F > 0.0\sigma(F)$ |
| 195 No. of reflections, working set | 28065 |
| 196 No. of reflections, test set | 1463 |
| 197 Final $R_{\text{crist}}$ | 0.163 |

|  |  |  |
| --- | --- | --- |
| 198 | Final $R_{\text{free}}$ | 0.195 |
| 199 | Cruickshank DPI | 0.08 |
| 200 | No. of non-H atoms: protein / water and solvent / total | 1844 / 168 / 2012 |
| 201 | R.m.s. Z deviations: bonds / angles | 0.85 / 0.88 |
| 202 | Average $B$ factors ($\text{\AA}^2$): protein / water and solvent / total | 30.0 / 36.5 / 30.5 |
| 203 | Ramachandran plot: favoured / allowed / outlier (%) | 98.8 / 1.2 / 0.0 |
| 204 | Molprobity score | 1.07 (99<sup>th</sup> percentile) |
| 205 | Molprobity clash score | 2.83 (99<sup>th</sup> percentile) |
| 206 | Rotamers: favoured / outlier (%) | 99.1 / 0.9 |

### 2.4. Structure solution and refinement

An initial single chain model was obtained by molecular replacement with *PHASER* 2.7 (McCoy *et al.*, 2007) using PDB entry 3ezj (Korotkov *et al.*, 2009) as template. Subsequently, automated building and refinement was performed by ARP/wARP (Langer *et al.*, 2008), which built 116 residues in chain A and 103 residues in chain B; and 6 additional residues that were not assigned to a chain, which were manually attached to the N-terminus of chain B. The model was further optimized using the PDB-REDO webserver (Joosten *et al.*, 2014), which also completed the missing loops (van Beusekom *et al.*, 2018). The optimal resolution cut-off was determined at 1.64 Å by PDB-REDO, which does a paired refinement as described by Karplus and Diederichs (2012).

Thereafter, the model was alternately refined with REFMAC5 (Murshudov *et al.*, 2011), manually rebuilt using COOT (Emsley *et al.*, 2010) and validated with *MolProbity* (Chen *et al.*, 2010). The N-terminus of chain B and the C-termini of both chains were extended by one residue, yielding a final total of 120 residues in chain B and 119 residues in chain A. The TLS groups were optimized to two groups per chain: one from the N-terminus up to residue number 90, and the second from 91 to the C-terminus. Side-chain alternates were modelled for 17 amino acids in total. Additionally, two main-chain alternates were built for the Gly29 and Gly30 of the first chain. Notably, the conserved cysteine bridge shows two clearly distinct conformations in chain B.

To analyse the conformation flexibility of the CDR loops, the structure was also refined with *phenix.emsemble_refinement* (Burnley *et al.*, 2012).

Molecular graphics figures were prepared with *CCP4mg* (McNicholas *et al.*, 2011).

## 3. Results and discussion

Nb6 binds very tightly to JBP1 (Fig. 1): the K_D_ was determined to be 30 nM. However, despite the high affinity for its target Nb6 unexpectedly crystallised without its intended binding partner JBP1, when set-up for crystallisation trials. It is therefore important to realize that despite very high binding affinity, separate crystallization of the nanobody is perfectly possible. The overall structure of Nb6 forms the expected immunoglobin fold of a nanobody (Fig. 2A) (Duhoo *et al.*, 2017). The three CDR domains are located at residues 48-56 (CDR1), 71-77 (CDR2), and 119-129 (CDR3). After superposition, the Cα RMSD between the two chains is 0.56 Å. The RMSD between the CDR1, CDR2, and CDR3 loops themselves in the different chains is only 0.41 Å, 0.27 Å, and 0.51 Å,respectively. Instead, a *β*-hairpin (residue 61-64) and some residues near the termini adopt distinct conformations.

**Figure 1.**
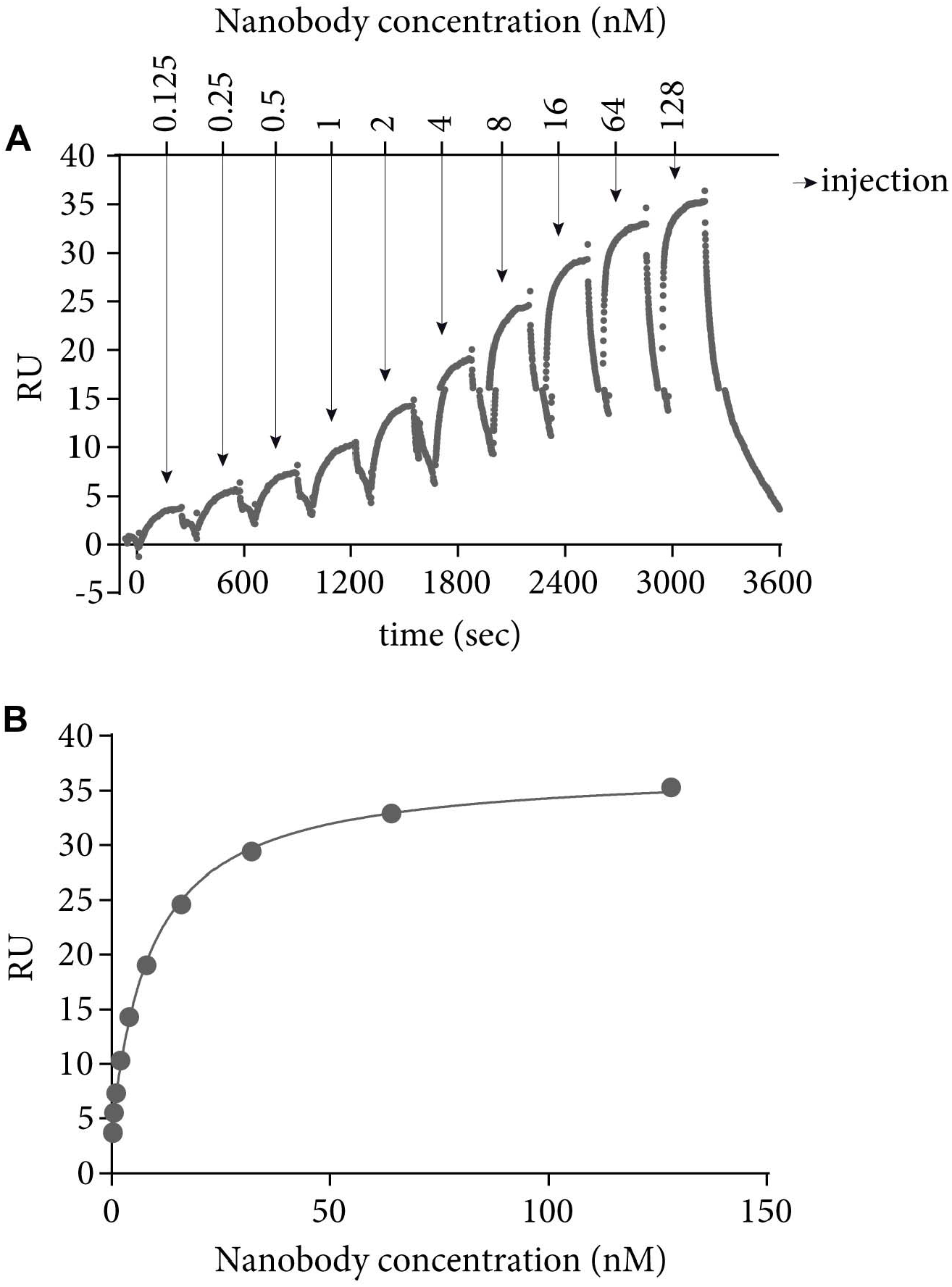
(A) A sensogram showing the substraction of the empty flow cell from the binding profile of the flow cell with immobilized JBP1. (B) The equilibrium response versus concentration. The equilibrium binding fit is shown as a black line.

**Figure 2.**
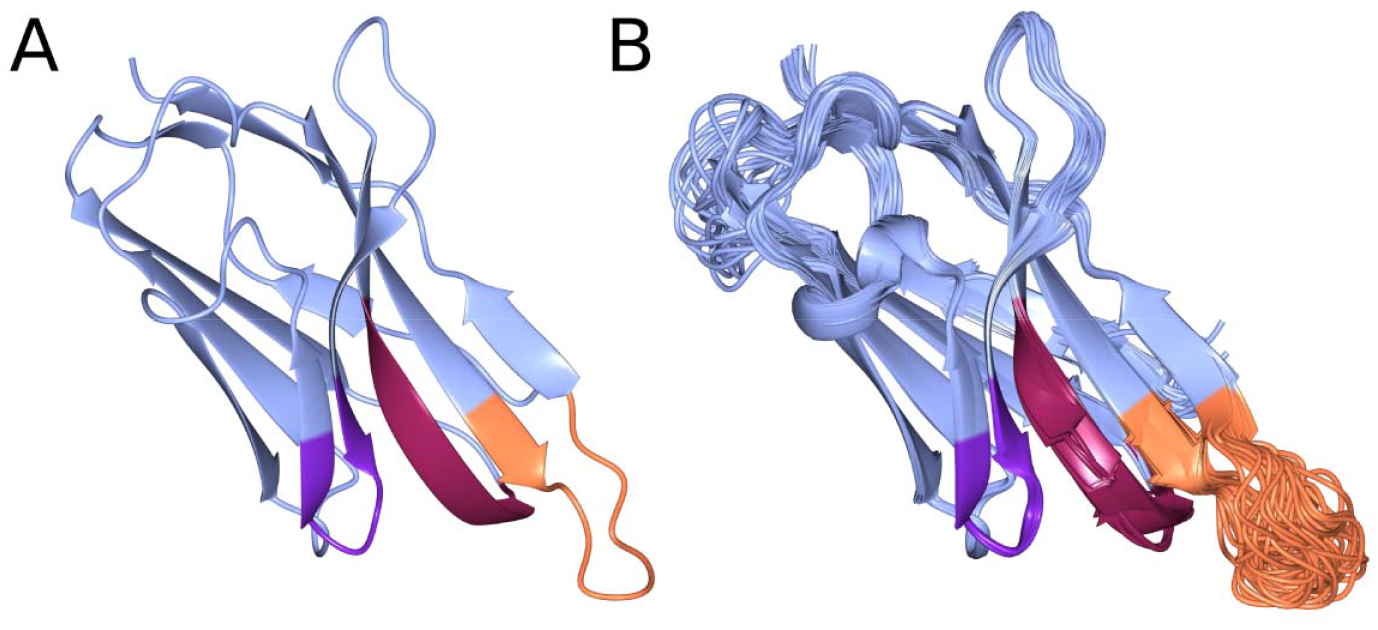
An overview of the nanobody structure model. (A) The nanobody structure model and (B) the same model after ensemble refinement. Chain A is shown in ice blue; chain B is shown in white. The CDR regions are shown in purple-red (CDR1), violet (CDR2), and orange (CDR3). From the ensemble refinement model, it is obvious that CDR1 and CDR2 are in a very fixed conformation, while CDR3 is much more flexible.

The average protein B factor was 30 Å^2^. Surprisingly, the average B factors for the CDR1 and CDR2 are mostly below average: in chain A and B, CDR1 has average B factors of 31 and 25 Å^2^, respectively. For CDR2, the average B-factor values were still lower at 21 and 19 Å^2^. CDR3 is much more flexible with average B factors of 47 and 58 Å^2^. The middle of CDR3 in chain B has the highest B factors in the entire structure model: average B factors of 96 and 110 Å^2^ were found for Tyr125 and Arg126, respectively. The flexibility of CDR3 was further confirmed by ensemble refinement (Fig. 2B), in which this loop adopts a wide range of different conformation. In contrast, CDR1 and CDR2 maintain largely the same conformation as in the single structure model. Furthermore, the electron density is very well-defined for CDR1 and CDR2, while density for CDR3 is only clear for the main chain at lower contour levels (Fig. 3).

**Figure 3.**
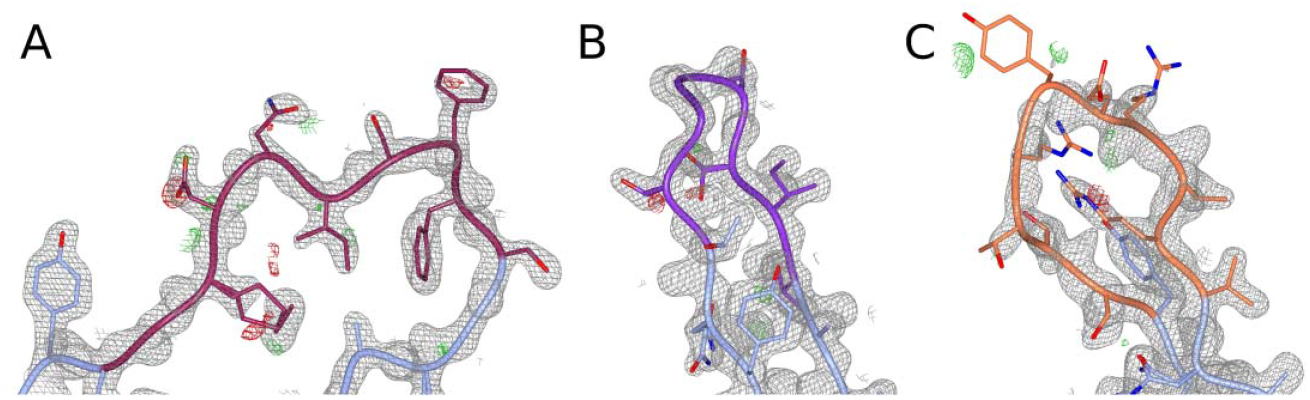
Details from the CDR regions of chain B: (A) CDR1, (B) CDR2 and (C·) CDR3. The 2 mF° - DF_c_ map is shown at 1.5 *σ* for CDR1 and CDR2, and at 1.0 *σ* for CDR3. The mF_o_ - DF_c_ map is shown at 3.0 *σ* in all cases. The backbone is shown as a tube for clarity; for side-chains, all atoms are shown. The density for CDR1 and CDR2 is very well-defined; for CDR3, however, the contour level had to be decreased to show clear density. Still, there is hardly any density observed for the side-chains in the tip of CDR3. Partly, this can be explained because there are two arginines and a glutamic acid in the tip, which are often flexible side-chains in general.

It should be noted that crystal contacts or non-crystallographic contacts may cause loops to appear more rigid than they are in solution, while intramolecular contacts such as interactions within the *β*-sheet are also present *in vivo.* In fact, apart from intramolecular contacts, all three CDR domains are also forming (non-)cyrstallographic contacts to some extent.

CDR1 forms a tight *β*-hairpin with the outer edges anchored in the *β*-sheet, which explains why it is ordered in general. In chain A, the tip is further stabilized by a hydrogen bond between the side-chain of Asn53 and a crystallographically related Gly63 backbone. This is not the case for chain B; however, there the first residues are stabilized further by strong *β*-sheet-like hydrogen bonding backbone of Phe49 with Ser38 from chain A.

The stabilization of of the first half of CDR2 is due to intramolecular *β*-sheet hydrogen bonding: the largest part of this variable domain is an extension of the conserved *β*-strand. In contrast, the last few residues form a *β*-hairpin and these residues form intermolecular contacts in the protein crystal. In both chains, extensive crystal contacts are formed for the last three residues in de CDR (residues 75-77), and the succeeding residues (78-80) even form a *β*-sheet with a crystal copy of Nb6.

The most flexible loop, CDR3, is far less influenced by crystal contacts. The tip of CDR3 forms some crystal contacts with its side-chains in chain A; however, these side-chains are not well-ordered so these contacts may not be so strong.

The analysis of RMSD, B factors, ensemble refinement and electron density consistently show that CDR1 and CDR2 have a very rigid conformation in Nb6. In contrast, CDR3 is much more flexible and there is only clear density for this loop at lower contour levels, implying that CDR3 only gets ordered upon binding to its target, JBP1. However, CDR1 and CDR2 are also involved in (non-)crystallographic contacts, which could cause them to appear more ordered than they are in solution. The extensive *β*-sheet interactions between different Nb6 monomers in the protein crystal may explain why it has crystallized in the absence of JBP1.

## Acknowledgements

The authors would like to thank Instruct-ERIC part of the European Strategy Forum on Research Infrastructures (ESFRI), and the Research Foundation Flanders (FWO) for their support to the Nanobody discovery. We thank Katleen Willibal for technical assistance during Nanobody discovery; we would also like to express our deep gratitude to the involved llamas.

